# Site specific target binding controls RNA cleavage efficiency by the Kaposi’s sarcoma-associated herpesvirus endonuclease SOX

**DOI:** 10.1101/320929

**Authors:** Aaron S. Mendez, Carolin Vogt, Jens Bohne, Britt A. Glaunsinger

## Abstract

During lytic replication of Kaposi’s sarcoma-associated herpesvirus (KSHV), the gene expression landscape of a cell is remodeled to evade the immune response and create an environment favorable to viral replication. A major driver of these gene expression changes is a virally encoded, messenger RNA (mRNA)-specific endonuclease termed SOX. SOX cleaves the majority of cytoplasmic mRNAs, but does so at specific internal sites loosely defined by a degenerate sequence motif. If and how RNA sequence directs SOX targeting remained unknown. To address these questions, we used recombinant, highly purified SOX endonuclease in a series of biochemical assays to reconstitute the cleavage reaction *in vitro* and gain significant insight into the biochemical mechanism of both SOX target recognition and endonucleolytic cleavage. Using this system, we determined that cut site specificity is preserved with purified SOX and a validated target RNA and thus does not require additional cellular cofactors. Furthermore, we showed that SOX displays robust, sequence-specific RNA binding to residues proximal to the cleavage site, which must be presented in a particular structural context. The strength of SOX binding dictates cleavage efficiency, providing an explanation for the breadth of target RNA susceptibility observed in cells.

**Significance Statement:** Kaposi’s sarcoma-associated herpesvirus (KSHV) is an oncogenic human virus that causes Kaposi’s sarcoma, primary effusion lymphoma, and multicentric Castleman disease. During viral replication, KSHV expresses an enzyme called SOX that cuts and inactivates the majority of cellular messenger RNAs, preventing their translation into proteins. Some mRNAs are efficiently cleaved by SOX, while others are poorly cleaved, but the mechanistic basis underlying this selectivity has remained largely unknown. Here, we reveal that the efficiency of RNA cleavage is heavily impacted by RNA sequences proximal to the cleavage site, which serve as a SOX binding platform. This helps explain both the range of RNA cleavage efficiency observed in SOX-expressing cells as well as the sequence specificity underlying SOX targeting.

## Introduction

Viral infection dramatically reshapes the gene expression landscape of the host cell. By changing overall messenger RNA (mRNA) abundance or translation, viruses can redirect host machinery towards viral gene expression while simultaneously dampening immune stimulatory signals (1–3). Suppression of host gene expression, termed host shutoff, can occur via a variety of mechanisms, but one common strategy is to accelerate degradation of mRNA (1, 2). This occurs during infection with DNA viruses such as alphaherpesviruses, gammaherpesvirues, and vaccinia virus, as well as with RNA viruses such as influenza A virus and SARS and MERS coronaviruses (1, 4, 5). In the majority of these cases, a viral factor promotes endonucleolytic cleavage of target mRNAs. This strategy bypasses the normally rate limiting steps of deadenylation and decapping to effect rapid mRNA degradation by host exonucleases (1).

Virally encoded host shutoff endonucleases are usually specific for mRNA, yet broad-acting in that they target the majority of the mRNA population. This is exemplified by herpesviral nucleases, including the SOX endonuclease encoded by Kaposi’s sarcoma-associated herpesvirus (KSHV), an oncogenic human gammaherpesvirus that causes Kaposi’s sarcoma and B cell lymphoproliferative diseases (6, 7). KSHV SOX is a member of the PD-(D/E)xK type II restriction endonuclease superfamily that possesses mechanistically distinct DNase and RNase activities (8–10). The RNase activity of the gammaherpesvirus SOX protein has been shown to play key roles in various aspects of the viral lifecycle, including immune evasion, cell type specific replication, and controlling the gene expression landscape of infected cells (11–14). However, the mechanism by which SOX targets mRNAs remains largely unknown.

Sequencing data indicate that within the mRNA pool there appears to be a range of SOX targeting efficiencies; some transcripts are efficiently cleaved in cells, while others are partially or fully refractory to cleavage (15–19). Additionally, SOX has been shown to cut within specific locations of mRNAs in cells, further emphasizing that there must be transcript features that confer selectivity (16, 20). Indeed, a transcriptome-wide cleavage analysis indicated that SOX targeting is directed by a relatively degenerate motif, often containing an unpaired polyadenosine stretch shortly upstream of the cleavage site, which is located in a loop structure (20). Cleavage within an unpaired loop was confirmed in a recent crystal structure of SOX with RNA, although additional contacts that could confer sequence specificity were not observed (21).

Thus, a major outstanding question is how RNA sequence and/or structure contribute to SOX target recognition. In this context, it is unclear how sequence features surrounding the RNA cleavage site might impact SOX targeting, for example by changing its affinity for a given RNA or the efficiency with which cleavage occurs. To address these questions, we sought to reconstitute the SOX cleavage reaction *in vitro* using purified components. Using an RNA substrate that is efficiently cleaved by SOX in cells, we revealed that specific RNA sequences within and outside of the cleavage site significantly contribute to SOX binding efficiency and target processing. In particular, we found that the polyadenosine stretch previously predicted to be important is critical for SOX binding, and we experimentally verified the importance of an open loop structure surrounding the cleavage site. Finally, we demonstrated that this *in vitro* system faithfully recapitulates the initial endonucleolytic cleavage event that is an essential component of mRNA target specificity *in vivo.* Collectively, our data reveal that specific sequence features potently impact SOX binding, and thus provide key insight into the breadth of SOX targeting efficiency observed across the transcriptome. More broadly, this information provides a framework for better understanding the target specificity of endonucleases, which play central roles in mammalian quality control processes and viral infection outcomes.

## Results

### KSHV SOX cleaves RNA substrates endonucleolytically as a monomer

In cells, the mRNA fragments resulting from the primary SOX endonucleolytic cleavage are predominantly cleared by the host 5’-3’ exonuclease XRN1, while *in vitro,* RNA fragments are rapidly degraded by 5’-3’ exonucleolytic activity intrinsic to purified SOX (9). Thus, it has been challenging to analyze the initial endonucleolytic cleavage event that is an essential component of mRNA target specificity *in vivo.* Here, we sought to develop a biochemical system to address these questions.

Our prior analysis of SOX targets in cells identified the human *LIMD1* mRNA, which codes for a protein essential for P body formation and integrity, as being highly susceptible to cleavage by SOX (20). The minimum sequence required to directly cut the putative cleavage site in *LIMD1* in cells was mapped to a 54 nucleotide segment (*LIMD1* 54), and we therefore chose this as our model substrate to study SOX targeting *in vitro* (20). We first expressed and purified KSHV SOX to greater then 95 percent purity from SF9 insect cells (fig. S1*A*). Using the *LIMD1* 54 substrate, we plotted the observed rate constant (*k*_obs_) as a function of SOX concentration, yielding a Hill coefficient of *n = 1.11* (Fig. 1*A*). Thus, in agreement with previous observations (9, 10), SOX appears to function predominantly as a monomer. Under conditions of half maximal activity (2 μM; Fig. 1*A*), SOX displayed a strong preference for the “hard” divalent metal Mg^2+^ and a weaker preference for the “softer” and larger metals Mn^2+^, Co^2+^, and Zn^2+^ (Fig. 1*B*). This is again consistent with other characterized members of the P/DExK family of enzymes (9, 22). Notably, SOX activity in the presence of Mg^2+^ was inhibited in a dose-dependent manner upon competitive addition of Ca^2+^ (Fig. 1*C* and S1*B*). This is likely the result of increased coordination partners engaged by Ca^2+^, which decreases the ability of catalytic residues to promote proper base hydrolysis (23–25). Finally, increasing the NaCl concentration above 100 mM led to substantially decreased SOX activity (Fig. 1D), in accordance with the observation that high salt concentrations frequently inhibit nuclease activity by disrupting protein-protein or protein-substrate interactions (25). Given that recombinant SOX displays robust 5’-3’ exonuclease activity (9, 10), we sought to confirm that *LIMD1* 54 was subject to endonucleolytic SOX cleavage, as this is the predominant event that directs mRNA turnover in SOX expressing cells (3, 16). Both the 5’ and 3’ ends *LIMD1* 54 were blocked by capping the 5’ end with a Cy5 fluorophore and the 3’ end with an Iowa Black quencher (*LIMD1* 54 Flo). We confirmed this RNA was resistant to degradation by the 5’-phosphate dependent exonuclease terminator (Fig. 1*E*, lane 3). However, in the presence of SOX, a cleavage product was observed that correlated with an endonucleolytic cut (Fig. 1*E*, lane 2).

**Fig. 1.**
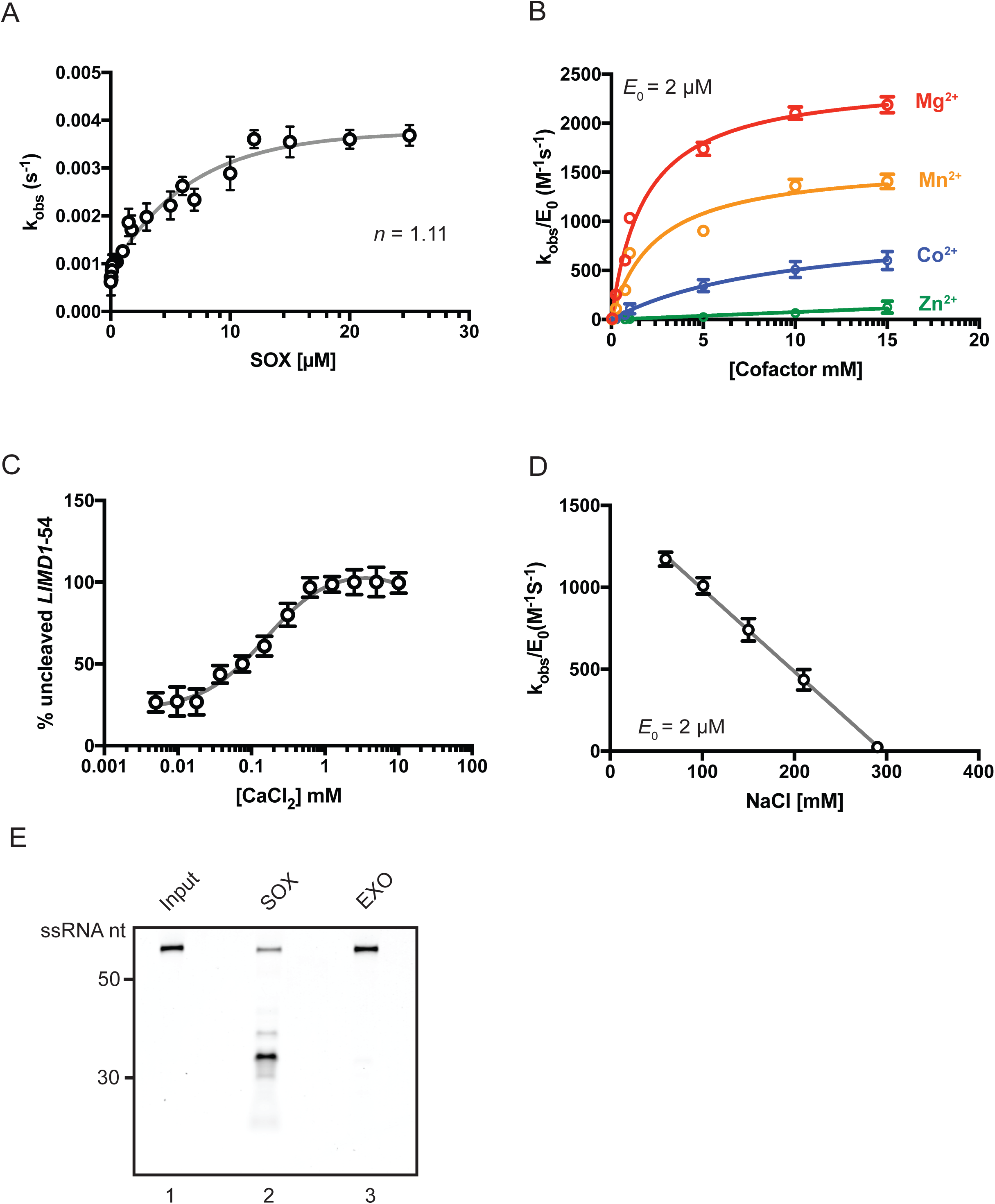
Kinetic characterization of recombinant SOX. (A) The observed rate constant (*k*_obs_) was plotted as a function of SOX concentration using the 5’ ^32^P-labeled *LIMD1 54* RNA substrate, showing a hill coefficient (*n*) of 1.11. (B) The catalytic efficiency of SOX (2μM) in the presence of MgCl_2_, MnCl_2_, CoCl_2_, and ZnCl_2_ was plotted as a function of cofactor concentration. (C) The impact of adding increasing concentrations of CaCl_2_ on SOX-induced degradation of a 5’ ^32^P-labeled *LIMD1 54* RNA probe. Reactions were carried out in the presence of 0.7 mM MgCl_2_. (D) SOX catalytic efficiency was determined under increasing concentrations of NaCl. (E) The 5’ and 3’ ends of *LIMD 54* were blocked with Cy5 and Iowa Black, respectively, to prevent exonucleolytic degradation. Reactions were incubated for 30 min in the presence of SOX or, as a control, the Terminator 5’ exonuclease (EXO). Input refers to RNA in reaction buffer without enzyme.

To confirm this processing event was not a result of contamination, we purified a SOX mutant containing mutations within two key residues of the SOX active site (D221N/E244Q). Incubation of this mutant with *LIMD1* 54 over the course of 1.5 h yielded no RNA cleavage (Fig. S1C). Thus, recombinant SOX appears to target *LIMD1* 54 for endonucleolytic cleavage *in vitro*, as has been observed for this substrate in cells.

### KSHV SOX shows RNA substrate selectivity *in vitro*

To analyze RNA substrate selectivity using our *in vitro* assay, we first compared SOX degradation of *LIMD1* 54 to a 51-nucleotide sequence of the mRNA encoding GFP (*GFP* 51). We have previously shown that *GFP* mRNA is cleaved by SOX in cells, and that *GFP* 51 is the minimal sequence required to elicit cleavage (15, 16). The cleavage sites for *LIMD1* 54 and *GFP 51* are predicted to occur in an open loop region (Fig. 2*A*, red arrow). Upon direct comparison of these two RNAs, we observed a 5-fold increase in the catalytic efficiency of SOX for the *LIMD1* 54 substrate compared to *GFP 51* (Fig. 2*B*). This difference was not exclusively due to the fact that the GFP substrate was slightly shorter than *LIMD1 54,* as SOX also displayed a 2.5-fold reduction of catalytic efficiency on a longer, 100 nt GFP substrate (*GFP* 100; Fig. 2*B*). Electrophoretic mobility shift assays (EMSA) further revealed a 10-fold increase in SOX binding to *LIMD1* 54 compared to *GFP* 51 (Fig. 2C). Given that both substrates contain the requisite unpaired bulge at the predicted cleavage site (see Fig. 2*A*), these observations suggest that additional sequence or structural features impact SOX targeting efficiency on individual RNAs.

**Fig. 2.**
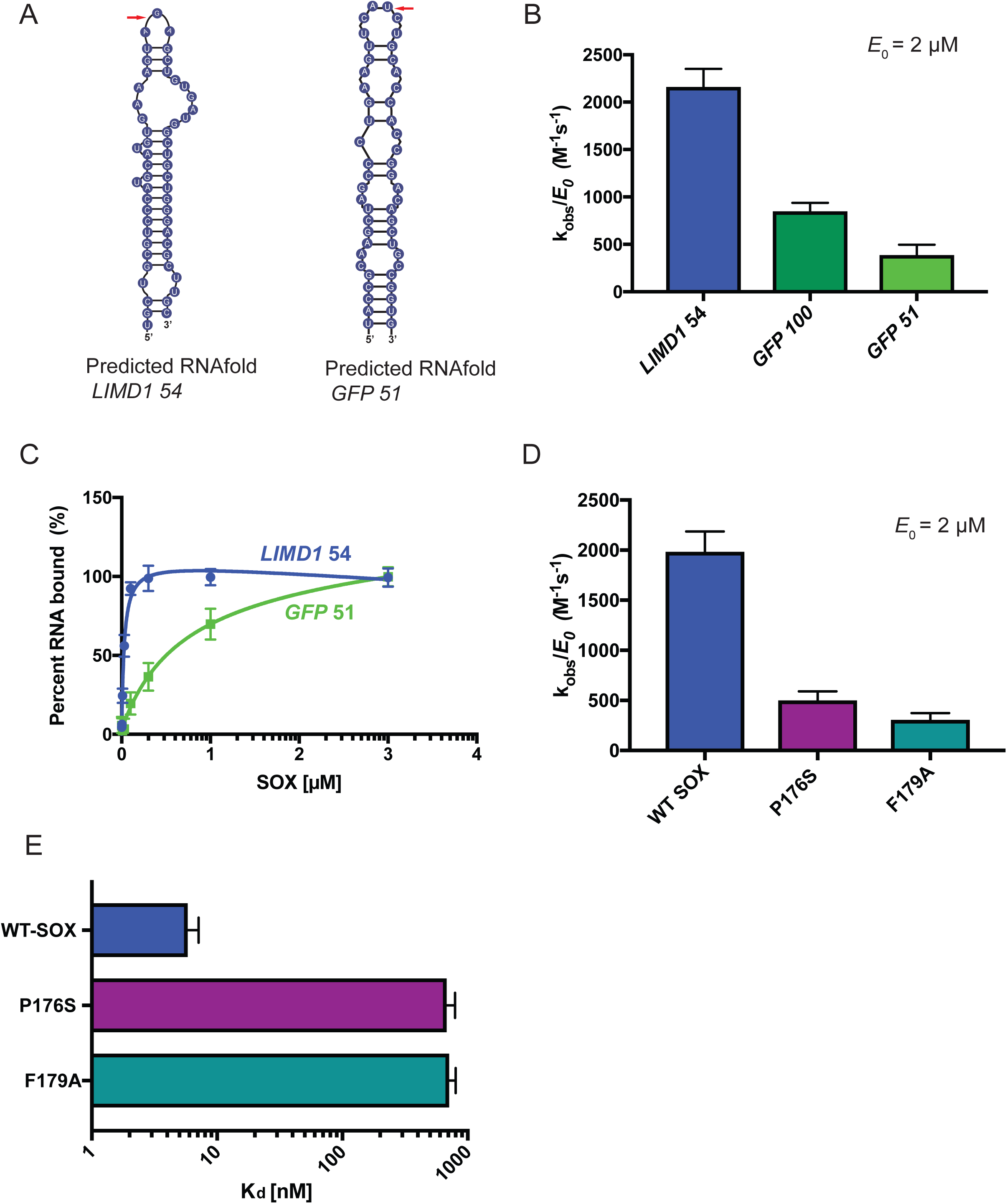
Substrate specificity, salt concentration and the bridge motif play important roles in SOX activity. (A) The predicted folding of the *LIMD1* 54 and *GFP* 51 RNAs was determined using mFold [Cite *Nucleic Acids Res.* **31 (13)**, 3406–15, (2003)]. Red arrows mark the predicted SOX cleavage site. (B) Catalytic efficiencies were determined for SOX (2 μM) in the presence of *GFP* 51, *GFP* 100 or *LIMD1* 54 substrate. Reactions were performed in triplicate. (C) Binding curves of SOX with *GFP* 51 and *LIMD1* 54 RNA. Percent binding of substrates was determined by EMSA, whereupon curves were fit to a single binding model from three independent measurements. (D) Catalytic efficiency of WT SOX or the host shutoff mutants P176S and F179A was determined at a constant enzyme concentration (2 μM) using a 5’ ^32^P-labeled *LIMD1 54* RNA probe. Experiments were performed in triplicate. (E) EMSAs were used to determine percent binding of 5’ ^32^P-labeled *LIMD1 54* RNA probe to WT SOX, P176S, and F179A. Curves were fit to a single binding model from three independent measurements.

Two SOX point mutants, P176S and F179A, located in an unstructured region of the protein that bridges domains I and II have been shown to be selectively required for its endonucleolytic processing of RNA substrates (Fig S2*A* and S2*B*) (8, 21). Structural data indicate that residue F179 forms a stacking interaction with an adenine base in the RNA, likely stabilizing the protein-RNA interaction, while P176 is hypothesized to contribute to structural rearrangements required for F179 engagement (21). We purified both mutants to evaluate their relative RNA processing and RNA binding activity against the optimal *LIMD1 54* substrate. Both mutants displayed purity and elution profiles similar to wild type (WT) SOX (see Fig. S1A). However, the catalytic efficiency of each mutant was >5-fold less than WT SOX (Fig. 2D). Furthermore, RNA binding was severely perturbed; the binding kinetics of WT SOX for *LIMD1* 54 are in the single digit nanomolar range (K_d_ = 7 nM), while P176S and F179A display >2 log defects (K_d_ = 702 nM and 831 nM, respectively) (Fig *2E* and S3*A-C*). Thus, the large defect in RNA binding likely explains the decreased efficiency of RNA processing. Notably, while there was a dramatic decrease in the relative affinities of the two mutants for *LIMD1* 54, there was not a complete loss of binding or RNA processing. This could be a result of secondary nonspecific interactions and/or nonspecific exonucleolytic degradation by SOX from the 5’ monophosphorylated end of the probe.

### Secondary structure determination of the *LIMD1* 54 substrate

*In silico* RNA folding predictions of SOX targeting motifs, coupled with RNA mutagenesis experiments, have indicated that an RNA stem loop structure is an important determinant in SOX targeting both *in vitro* and *in vivo* (20, 21). Given the importance of this predicted motif, and in particular the proposed requirement for unpaired sequence at the cut site, we sought to experimentally determine the structure of *LIMD1* 54 using chemical based in-line probing (Fig. 3A). This showed that the *LIMD1* 54 structure contains a largely base paired stem region, followed by a loop at positions 15–27 that encompasses the predicted SOX cleavage site between nt 26 and 27, and a short hairpin structure at positions 29–40 (Fig. 3B). Notably, some differences exist between the predicted and observed structures of *LIMD1* 54, including a larger loop region and the subsequent short stem-loop (compare Fig. 3B to Fig. 2A) However, in both cases the predicted cleavage site of SOX resides in a loop region.

**Fig. 3.**
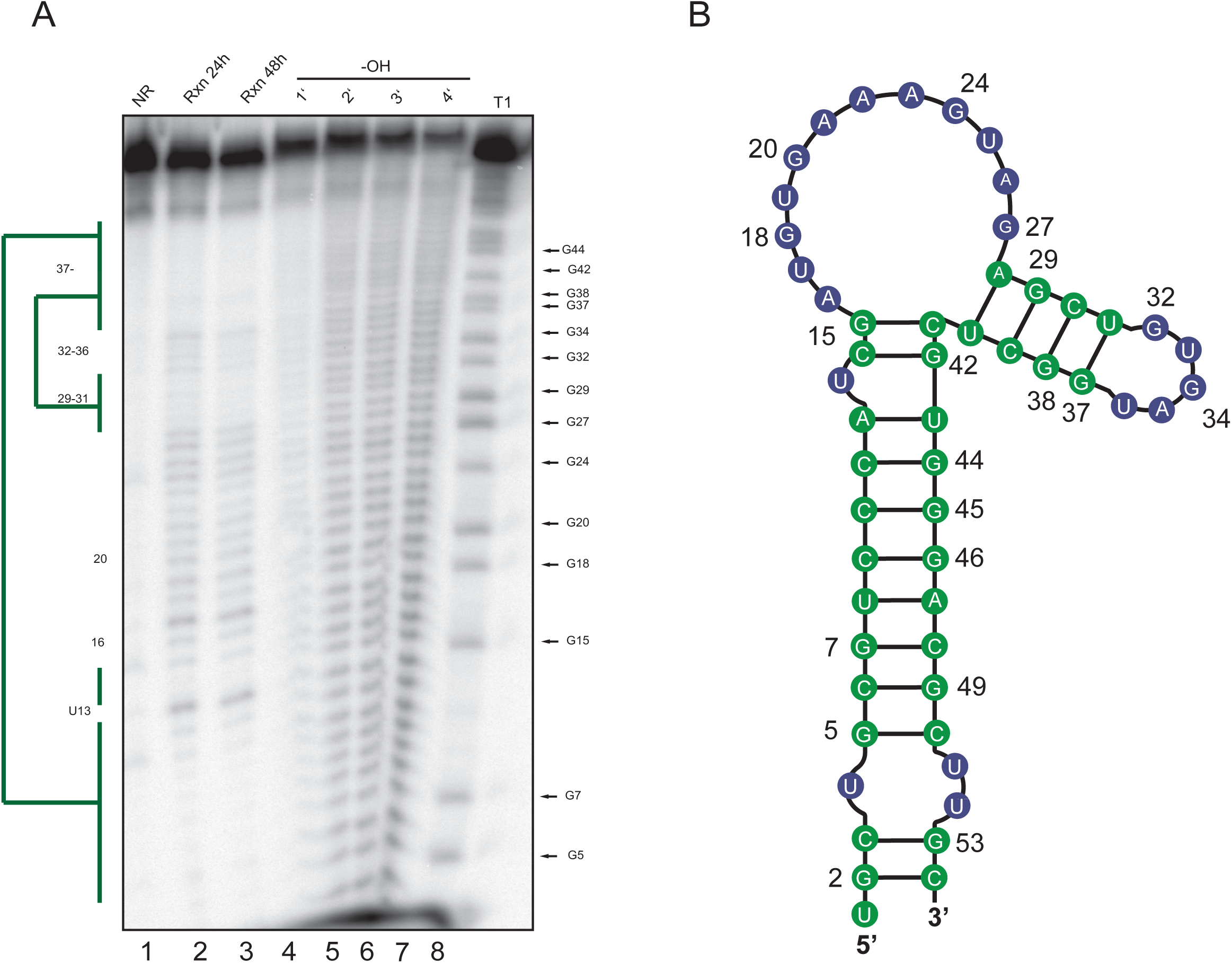
In-line structure probing of *LIMD1* 54. (A) An in-line reaction (Rxn) was performed at room temperature for 24 h (lane 2) or 48 h (lane 3) at pH 8.3 to identify structured regions of the *LIMD1* 54 RNA. Ladders were generated by subjecting the RNA to cleavage by RNase T1 (lane 8) or alkaline hydrolysis (-OH, lane 4–7). Products were separated by 8% urea PAGE, whereupon structured regions protected from cleavage were identified (green bars). No reaction (NR, lane1) refers to the input RNA. (B) Diagram showing the *LIMD1 54* structure as deduced from the in-line probing gel. Green color corresponds to the structured regions denoted by the bars in (A), while blue refers to unstructured regions.

### SOX binds to an unpaired stretch of adenosine repeats

Recently, a high-resolution crystal structure was solved of SOX bound to a 31nt fragment of the KSHV pre-microRNA K12–2 (K2–31). In this structure, the only observed contacts between SOX and K2–31 occurred between the four active site residues of SOX (Y373, R248, C247, F179) and the UGAAG motif surrounding the cleavage site of the RNA (21). It was therefore hypothesized that no other residues beyond this unpaired UGAAG motif were involved in transcript recognition (21). However, our data showing a 10-fold difference in binding affinity between two *in vivo* validated SOX substrates suggested that a more extended interaction surface might distinguish optimal from sub-optimal RNA substrates. We therefore used RNA footprinting to map the SOX binding sites on *LIMD1 54.* Indeed, SOX robustly protected 3 *LIMD1 54* adenosines (positions 20–24) from RNase I digestion in a dose dependent manner (Fig. 4). Notably, this mapped binding site is the same one predicted from *in vivo* PARE-seq data (20). We also observed a modest protection of base 27 (G) located directly adjacent to the predicted cleavage site of SOX, which represents the region detected in the crystal structure of K2–31 bound to SOX. Collectively, these findings suggest that while SOX may interact with residues directly adjacent to the cut site, a more extensive interaction interface exists for its preferred *in vivo* targets.

**Fig. 4.**
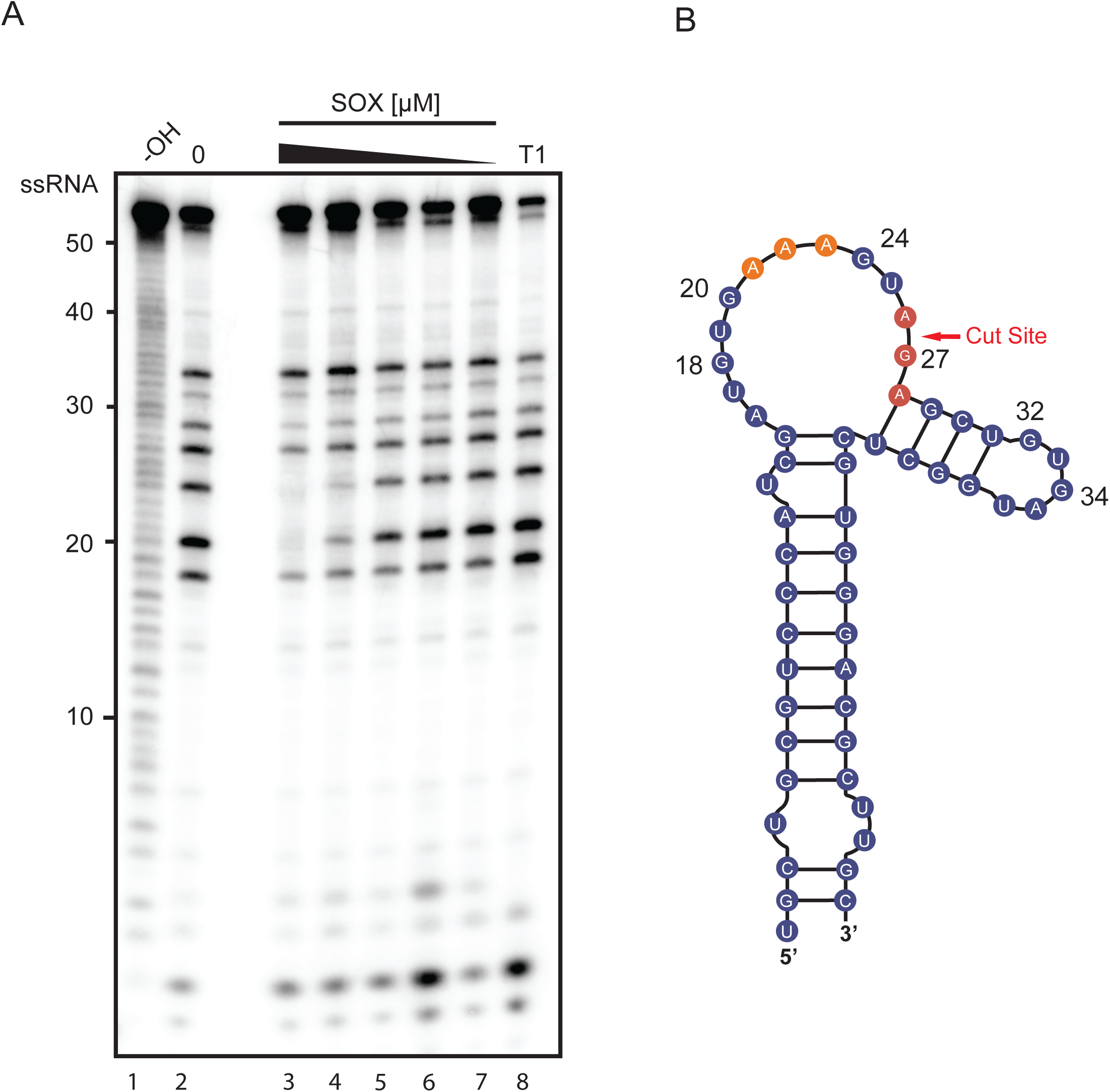
SOX binds to a stretch of adenosines upstream of the cleavage site. (A) An RNA footprinting assay was carried out by incubating 5’ ^32^P-labeled *LIMD1 54* with RNase I in the presence (lanes 3–7) or absence (lane 2) of a dilution series of SOX (8–0.5 μM). Hydrolysis (-OH, lane 1) and RNase T1 (T1, lane 8) ladders of the RNA were also generated in order to map the location of protected sites. (B) Diagram of *LIMD1* 54 indicating sites protected from RNase I cleavage by SOX. The upstream SOX binding site is colored orange while the protected residues surrounding the cut site are shown in red.

### Residues surrounding the SOX binding and cleavage sites contribute to efficient substrate degradation

To explore the importance of the residues involved in SOX binding and cleavage, we engineered 3 mutants of the *LIMD1 54* substrate (Fig. 5A). First, we preserved the loop structure but replaced the 3 adenosines bound by SOX (residues 39–43) with guanosines (*LIMD1* 3xA-G). Second, we mutated the residue located at the predicted SOX cut site that was also protected in the footprinting assay (*LIMD1* A-G). This mutant has been previously identified to block SOX cleavage *in vivo* (20). Third, we largely abolished the loop structure by providing complementary base pairing (*LIMD1* Zipper). The predicted structures of the *LIMD1* 3XA-G and *LIMD1* zipper mutants were verified by in-line probing (Fig. 5, S6). Real-time binding kinetics for SOX with WT *LIMD1* 54 and each of the 3 mutant substrates were then measured using bio-layer interferometry (BLI). All RNA probes were 3’ biotinylated and immobilized to a streptavidin-coated BLI probe, whereupon the binding and dissociation of SOX was measured. To prevent degradation of the probe, excess calcium ion was used in place of magnesium (Fig. S1B). SOX retained similar binding affinity to the cut site mutant *LIMD1* A-G (K_d_ = 25 nM) as to WT *LIMD1* 54 (K_d_ = 16.3 nM) (Fig. 5B, S4A-B and table S1). In contrast, SOX exhibited dramatically reduced binding to both the predicted binding site mutant *LIMD1* 3xA-G (K_d_ = 710 nM) and to the *LIMD1* zipper mutant lacking the loop region (K_d_ = 904 nM) (Fig 5B, S4C-D and table S1). We also measured SOX binding to the KSHV pre-miRNA sequence used to obtain the SOX-RNA co-crystal structure (K2–31) (21). Notably, the affinity of SOX for K2–31 was even less than that of the *LIMD1* 3xA-G and Zipper mutants (K_d_ = 1.8 μM), suggesting that despite having an UGAAG motif upstream of a predicted bulge, this may not be a bona fide SOX target (Fig. 5*B*, S4*E* and table S1).

**Fig. 5.**
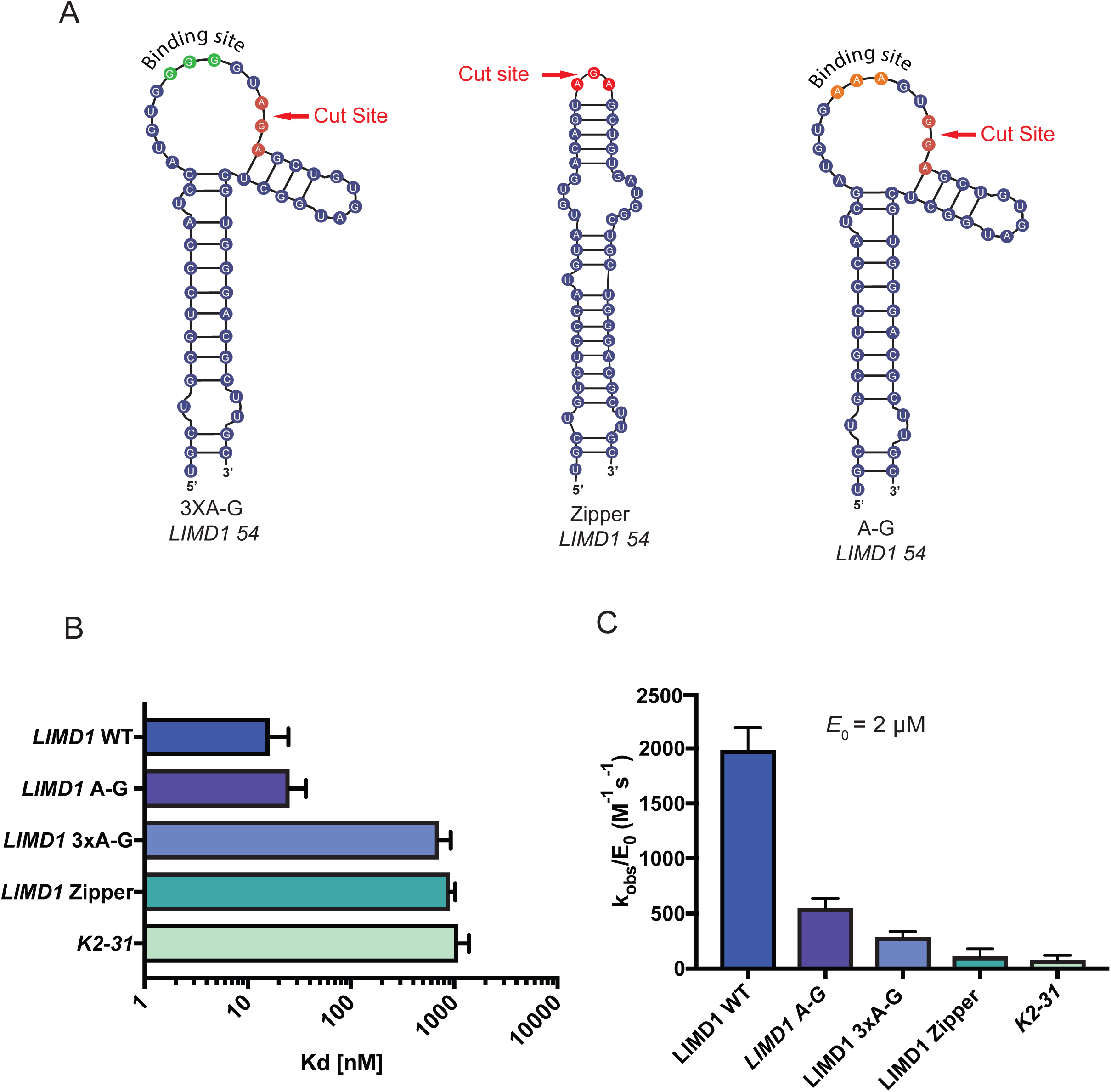
Mutations that disrupt *LIMD1* 54 structure reduce SOX binding affinity. (A) Diagram showing the location of mutations made within *LIMD1* 54. Binding site mutants and cut site mutants are labeled accordingly. (B) The binding affinity of SOX for each of the *LIMD1* 54 mutants was tested in parallel using Bio-Layer Interferometry (BLI). (C) SOX catalytic efficiency was tested at a constant concentration (2 μM) for all of the *LIMD1* 54 mutant substrates. All assays were performed in triplicate.

We next quantitatively measured the catalytic efficiency of SOX towards each of the above RNA substrates. Despite SOX having WT binding affinity for the predicted cleavage site mutant *LIMD1* A-G, there was a 4-fold defect in its ability to degrade this mutant (Fig. 5C). Even more marked defects in SOX catalytic efficiency were observed for the binding site mutant *LIMD1* 3XA-G, the loop mutant *LIMD1* zipper, and the pre-miRNA K2–31 (Fig. 5C). Collectively, these data indicate that efficient RNA cleavage requires both an appropriate SOX binding site and a suitable cut site.

### Site-specific endonucleolytic cleavage of target RNA occurs in vitro

In cells, SOX cleaves its mRNA substrates site-specifically. Mutagenesis of residues in mapped cleavage sites generally abolishes SOX cleavage at that location (20). To determine if our *in vitro* assay faithfully recapitulated the site specificity of SOX endonucleolytic targeting observed in cells, we established reaction conditions that enabled trapping of the early cleavage events. By combining Ca^2+^ and Mg^2+^ in our reaction buffer, we were able to sufficiently slow SOX processing to visualize cleavage products derived from 5’ ^32^P labeled substrates. Indeed, we observed a predominant 27 nt band, which is the size of the product released upon *LIMD1 54* cleavage at the predicted cut site (Fig. 6*A*, lane 3). Additional bands also appeared, likely representing subsequent processing events. Importantly, when we incubate SOX with the cut site mutant *LIMD1 A-G,* there is a complete loss of this 27 nt product, as well as the additionally processed intermediates (Fig. 6A, lane 4). Production of these cleavage intermediates required SOX, as no decay was observed in the RNA-only controls (Fig. 6A, lanes 1–2).

**Fig. 6.**
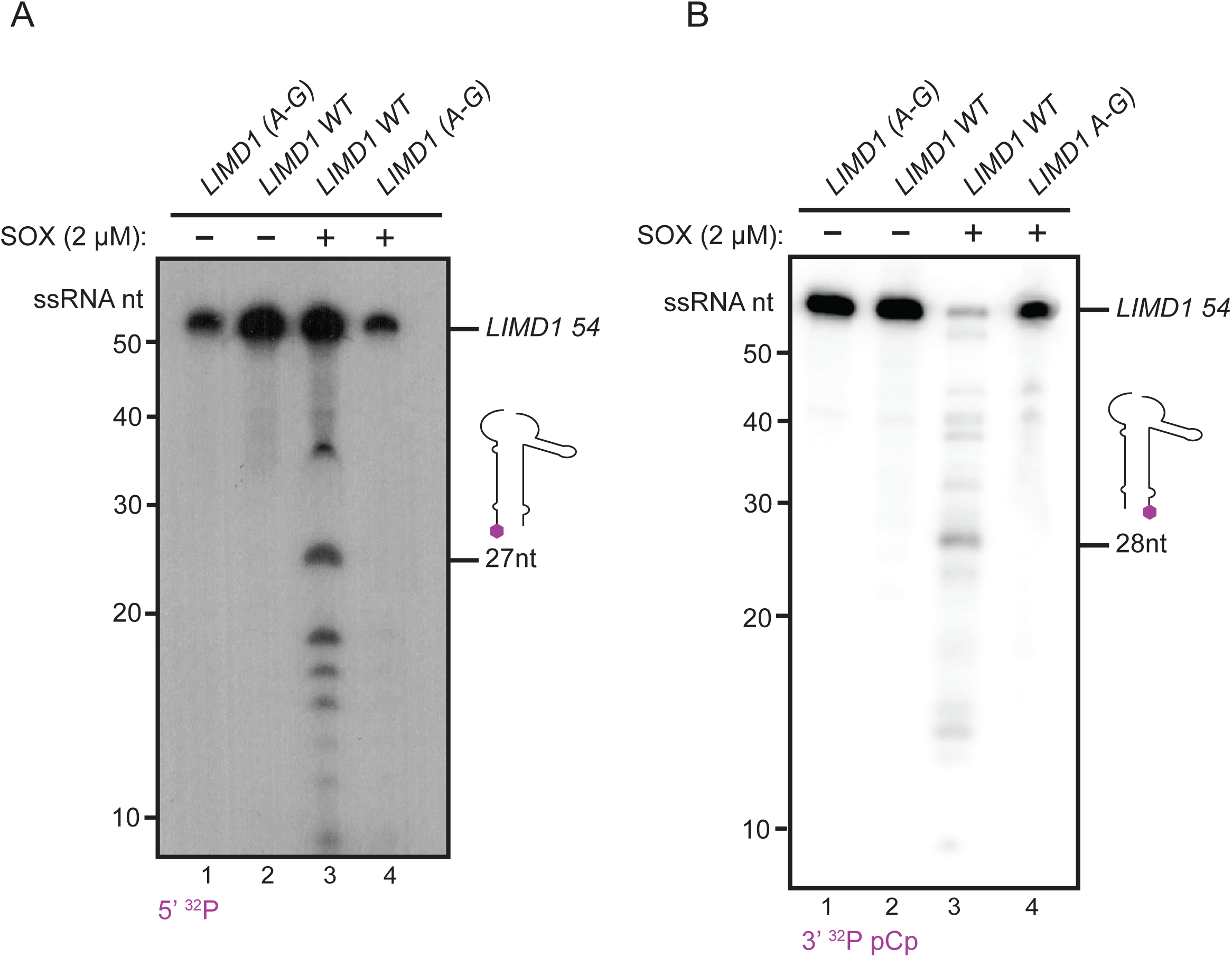
SOX endonucleolytically cleaves *LIMD1* 54 in a site-specific manner. (A) 5’ ^32^P-labeled *LIMD1 54* and *LIMD1 54* A_G were incubated for 10 min in 0.1 mM CaCl_2_ and 0.7 mM MgCl_2_ in the presence or absence of 2 μM SOX. The 5’ cleavage product with a size corresponding to cleavage at the mapped cut site (27 nt) is shown. (B) Cleavage assay was performed as in (A), except using 3’ ^32^P pCp-labeled *LIMD1* 54. The 3’ cleavage product with a size corresponding to cleavage at the mapped cut site (28 nt) is shown.

Finally, we sought to verify that the predominant 27 nt cleavage product we observed was a result of an endonucleolytic cleavage and not 5’ end processing. To this end, we generated a *LIMD1* 54 substrate containing a 3’ ^32^P pCp label and a free 5’ OH to block 5’ end processing. Again, in the presence of SOX, WT *LIMD1 54* but not the A-G mutant produced a cleavage product whose size corresponded to cleavage at the predicted site (Fig 6*B*). Taken together, these data confirm that our *in vitro* assay faithfully recapitulates SOX cleavage site specificity on a true substrate.

## Discussion

Endonuclease-directed mRNA degradation plays key roles in the lifecycle of gammaherpesviruses, yet the fundamental principles governing target specificity by SOX are not well understood. Here, through the development of the first biochemical system to faithfully recapitulate the internal cleavage specificity observed for SOX in cells, we revealed how both RNA sequence and structure contribute to targeting. These findings resolve a central feature of the current model of SOX activity (Figure 7). Previous observations established that sequences flanking the cut site were required to direct cleavage by SOX (16, 20). However, it was unresolved whether they played a strictly structural role in presenting an exposed loop for cleavage, served as a platform for SOX binding, or created a binding site for one or more cellular factors that then indirectly recruited SOX to its targets. Through a combination of mutational analyses, RNA structure probing, and RNA footprinting assays, we showed that efficient SOX targeting requires both an exposed loop structure and upstream sequences that serve as a SOX binding platform. This combination of sequence and structural features within the targeting motif helps explain why some mRNAs are efficiently cleaved by SOX, whereas others are weaker substrates.

**Fig. 7.**
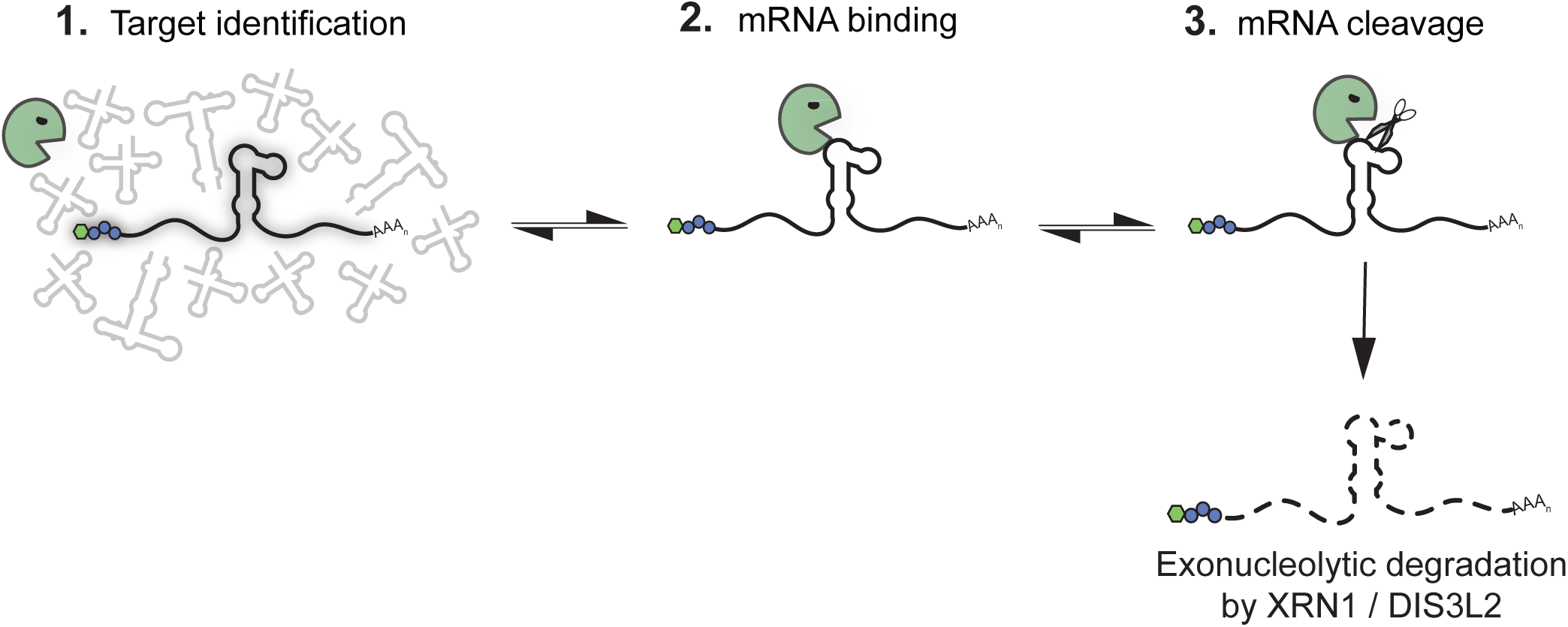
Model of mRNA targeting by SOX. SOX is able to distinguish mRNA from other types of RNA in cells by an as yet unknown mechanism. Subsequently, it endonucleolytically cleaves its targets at specific sites, whereupon the fragments are degraded by host exonucleases such as Xrn1 and Dis3L2. Here, we revealed that in addition to the requirement for an unpaired loop at the cleavage site, additional upstream RNA sequences increase the affinity of SOX for individual targets, thereby controlling cleavage efficiency.

A key open question related to SOX function is how it can target the majority of mRNAs in cells, yet with significant site specificity. These observations suggest that there must be specific mRNA features that influence targeting. Indeed, PARE-seq analyses of cleavage intermediates in SOX expressing cells revealed that cleavage sites were associated with a degenerate sequence motif (20). Sequences proximal to the cleavage site were predicted to be un-base paired and frequently contained a polyadenosine stretch followed by a purine (20). The importance of these sequence features for SOX targeting was validated for the *LIMD1* transcript in cells (20). Because *LIMD1* has been established as a particularly robust SOX target in cells (20), we reasoned that it must contain features optimal for SOX processing and therefore would be an ideal substrate to dissect these features biochemically. Indeed, SOX binding to *LIMD1* 54 was 10-fold better than to the commonly used reporter substrate GFP, and an order of magnitude better than to the K2–31 pre-miRNA, which has not been demonstrated to be processed by SOX in cells. Importantly, these binding differences correlated with the efficiency of SOX cleavage *in vitro,* arguing that the ability to bind the targeting motif is a key step in target recognition. Through RNA footprinting assays, we were able to show that SOX binds to a bulge structure proximal to the cleavage site containing the polyadenosine stretch previously predicted to be important (20). Mutating either the bulge structure (*LIMD1* zipper) or maintaining the bulge but mutating the polyadenosine stretch (*LIMD1* 3xA-G) resulted in an order of magnitude reduction in binding affinity, correlating with a dramatic decrease in cleavage efficiency. Collectively, these data demonstrate that variability in the efficiency of SOX targeting observed in cells is likely due to differences in RNA sequences that mediate SOX binding.

A recent crystal structure of SOX bound to the K2–31 pre-miRNA captured the importance of the exposed loop region for SOX cleavage (21). However, the structure did not reveal additional interactions between SOX and the RNA beyond the 3 residues surrounding the cut site. Our data suggest that this is likely because the K2–31 RNA lacks the additional residues necessary for SOX binding site found in both *LIMD1* and *GFP.* While the K2–31 RNA does contain adenosines upstream of the cleavage site, structural predictions indicate these residues are within a stem region (21), rather than in an exposed loop as is the case for *LIMD1* and *GFP.* Together, these observations indicate that while upstream adenosines are important for binding, they must be present in an unpaired state to promote SOX binding. It is notable that prior studies reported much weaker interactions between SOX and RNA (K_d_ = 75 μM) compared to its DNA substrates (K_d_ = 1 μm) (9, 10, 21). However, in these cases binding assays were conducted with scrambled RNA sequences. We found that SOX binding affinities to RNA substrates vary over several orders of magnitude, in a manner that correlates with cleavage efficiency. Interestingly, the crystal structure of SOX bound to DNA showed more dynamic interactions along the length of the protein (~480 Å^2^ interaction surface), when compared to the K2–31 RNA bound structure (~240 Å^2^ interaction surface). It is therefore possible that more interaction along the length of SOX protein might occur with optimal substrates such as *LIMD1* that are more tightly bound.

The fact that purified SOX endonucleolytically cleaved *LIMD1* 54 at the precise site observed in SOX-expressing cells demonstrates that cleavage site selection on an mRNA is not mediated by a cellular cofactor. Instead, targeting at particular RNA motifs is strongly influenced by the strength of SOX binding. Our observation that the P176S and F179A SOX mutants display significant RNA binding defects indicates that their failure to cleave mRNAs in cells is due to an inability to efficiently bind the targeting motif.

The mechanism by which SOX initially distinguishes RNA polymerase II transcribed mRNAs from other types of RNA in cells remains an important open question, as this feature of SOX selectivity is not preserved *in vitro.* We hypothesize that cellular co-factors, perhaps though interactions with SOX, enable this distinction. More broadly, endonucleases are instrumental in RNA processing and degradation. Nuclease processing defects lead to several human pathologies ranging from cancer to neurodegeneration (26–30), and our study provides a framework for better understanding the mechanistic features governing endonuclease targeting.

## Materials and Methods

Full details are available in *SI Materials and Methods*

### Recombinant protein expression and purification

KSHV SOX was codon optimized for Sf9 expression and synthesized from GENEWIZ. SOX was then subcloned using restriction sites BamHI and SalI (New England BioLabs) into pFastBac HTD. This vector was modified to carry a GST affinity tag and PreScission protease cut site as described (31). All SOX mutants were generated using single primer site-directed mutagenesis (32). Sequences were validated using standard pGEX forward and reverse primers. Generation of viral bacmids and transfections were prepared as described in the Bac-to-Bac^®^ Baculovirus Expression System (Thermo Fisher Scientific) manual. After transfection, Sf9 cells (Thermo Fisher Scientific) were grown for 96 hr at 22 °C using SF-900 SMF media (Gibco) substituted with 5% fetal bovine serum (FBS) and 1% antibiotic antimycotic (AA). Supernatant was transferred to a 6-well tissue culture plate containing 1 ml of 2×10^^6^ cells/well. Cells were incubated for 96hr to generate passage 1 (P1). The P1 supernatant was transferred to a flask containing 50 mL of 1×10^^6^ cells/ml and incubated for 96 h, a time point sufficient to yield 5 mg of SOX per 50 mL of cells. Protein expression was confirmed by Western blot with an anti-GST antibody (GE Health Care Life Sciences).

Sf9 cell pellets were suspended in lysis buffer containing 600 mM NaCl, 5% glycerol, 0.5% Triton X-100, 3 mM DTT, 20 mM HEPES pH 7.0 with a cOmplete, EDTA-Free protease inhibitor Cocktail tablet (Roche). Cells were sonicated on ice using a macro trip for 30 sec bursts with 1 min rests for 12 min at 50 Amps. Cell lysate was cleared using a pre-chilled (4°C) Sorvall LYNX 6000 Superspeed Centrifuge spun at 18,000 rpm for 45 min. The cleared lysate was incubated for 4 hr at 4 °C with rotation with 4 mL of a GST bead slurry (GE Healthcare Life Sciences) that had been pre-washed 3x with wash buffer (WB) containing 600 mM NaCl, 5% glycerol, 3 mM DTT, 20 mM HEPES pH 7.0. The bead-protein mixture was washed 3X times with 15 mL of WB, then transferred to a 10 ml disposable column (Qiagen) and washed with an additional 50 mL of WB followed by 100 mL of low salt buffer (LSB) containing 250 mM NaCl, 5% glycerol, 3 mM DTT, 20 mM HEPES pH 7.0 with periodic resuspension to prevent compaction. SOX was then cleaved on column with PreScission protease (GE Healthcare Life Sciences) overnight at 4°C, and remaining protein was collected with a final 8 mL LSB wash.

Cleaved protein was concentrated to ~ 1 mL using Amicon filter concentrator membrane cut off 30 kDa (EMD Millipore), then loaded onto a HiLoad Superdex S200 pg gel filtration column (GE Healthcare Life Sciences). Protein elutions were concentrated using an Amicon concentrator described above to 5 mg/mL and aliquots were snap frozen in liquid N_2_ and stored at − 80 °C.

### RNA substrate preparation and end labeling

All RNA substrates (sequences in Table S2) unless stated otherwise were synthesized by Dharmacon (GE Healthcare) with HPLC and page purification. RNAs were 5’ end labeled with γ-[^32^P]-ATP-6000 Ci/mmol 150 mCi/ml (Perkin Elmer) using T4 PNK (New England BioLabs). RNAs were 3’ end labeled with 5’-[^32^P]-pCp 3000 Ci/mMol 10 mCi/ml using T4 RNA ligase 1 (New England BioLabs). Labeled RNA substrates were purified using 20% urea-PAGE and were isolated from gel slices by incubating overnight at 8°C in a buffer containing 10 mM Tris-HCl, 1 mM EDTA pH 8.0. Eluted RNAs were ethanol precipitated and resuspended in RNase-free ddH_2_O.

### Ribonuclease assays

*k*_obs_ and Hill coefficients of SOX were determined from the cleavage kinetics of [^32^P]-labeled RNA substrates as previously described (32). Briefly, 1 μl (≤1 pM) of [^32^P]-labeled RNA was added to 9 μl of premixture containing 20 mM Hepes pH 7.1, 70 mM NaCl, 2 mM MgCl_2_, 1 mM TCEP, 1% glycerol, and increasing concentrations of purified SOX. Reactions were performed at room temperature under single turnover conditions, and quenched at the indicated time intervals with 8 μl stop solution (10 M urea, 0.1% SDS, 0.1 mM EDTA, 0.05% xylene cyanol, 0.05% bromophenol blue). Samples were resolved by 15% urea-PAGE, imaged using a Typhoon variable mode imager (GE Healthcare), and quantified using ImageQuant and GelQuant software packages (Molecular Devices). The data were plotted and fit to exponential curves using Prism 7 software package (GraphPad) to determine observed rate constants.

A FRET probe with excitation at 646 nM and emission at 662 nM (*LIMD1* 54 Flo) was purchased from Dharmacon. The RNA FRET probe was added at a final concentration of 100 nM to 9 μl of premixture containing 20 mM Hepes pH 7.1, 70 mM NaCl, 2 mM MgCl_2_, 1 mM TCEP, 1% glycerol with 2 μM of SOX (32). Terminator 5’ exonuclease (Lucigen) was added to reactions using a 1:1000 dilution of the enzyme. reactions were quenched at indicated time intervals with equal volumes of stop solution containing 95 *%* formamide and 10 mM EDTA, then resolved using urea-PAGE and visualized using a Typhoon variable mode imager (GE Halthcare). The data were plotted using Prism 7 software package (GraphPad). All experiments were repeated >3 times and mean values were computed.

For assays designed to detect endonucleolytic cleavage intermediates, 1 μl of labeled RNA substrate was combined with 9 μl of reaction solution (20 mM HEPES pH 7.1, 70mM NaCl, 0.200 mM CaCl, 0.700 mM MgCl_2_, 1 % glycerol, 0.5 mM TCEP) in the presence or absence of 2 μM SOX for 10 min at room temperature. RNA was then ethanol precipitated, resuspended in 95 % formamide solution containing 10 mM EDTA, and resolved on a 12% urea-PAGE analytical grade sequencing gel together with a ssRNA Decade ladder (Ambion Life Technologies) for 1.5 hours at 22 watts before imaging as described above.

## Acknowledgements

We thank members of the Glaunsinger lab for their suggestions and critical reading of the manuscript. This work was supported by NIH R01CA136367 (B.A.G.), C.V. was supported by the DAAD P.R.I.M.E. program (57222872) for postdoctoral fellows. B.A.G. is an investigator of the Howard Hughes Medical Institute.

## Supplementary Information

### SI Materials and Methods

#### In-line probing

The sequence surrounding the cut site in LIMD1 was inserted into a pBSSK (-) backbone using the BamHI and XbaI restriction sites. Mutations were introduced by the Quickchange site directed mutagenesis protocol (Agilent). The 100 nt sequence surrounding the GFP cut site was inserted using the BamHI and XhoI restriction site. In-line probing was performed as described previously (1). Briefly, pBSSK(-) plasmids containing the indicated sequences (see Table S2) were linearized by digestion with XhoI and ScaI for GFP or BlpI and SacI (NEB) for LIMD1, gel purified, phenol/chloroform extracted, and ethanol precipitated. The fragments were then used as templates for *in vitro* transcription with the HiScribe T7 High Yield RNA synthesis Kit (NEB) and afterwards subjected to Turbo DNase (Ambion by Life Technologies) treatment. RNA was revolved by 8% Urea PAGE, and full length transcripts were excised from the SYBR Gold stained gel (Thermo Fisher Scientific), eluted overnight in G50 buffer (20 mM Tris HCl pH 7.5, 300 mM NaOAc, 2 mM EDTA, 0.2% SDS), phenol/chloroform extracted, and ethanol precipitated. The RNA (~40 pmol) was dephosphorylated using shrimp alkaline phosphatase (rSAP, NEB), labeled with 1 μl [_Y_^32^P] ATP (150 mCi/ml) using USB Optikinase (Affymetrix), then gel purified as described above and dissolved in 20 μl of nuclease free water. For the in-line probing reaction, 1 μl RNA (≥ 20,000 cpm) was incubated in 2x reaction buffer (100 mM Tris-HCl pH 8.3, 40 mM MgCl_2_, 200 mM KCl) at room temperature for 24 h or 48 h. The reaction was quenched with 2x loading buffer (10 M urea, 1.5 mM EDTA pH 8.0). To generate ladders, 1 μl of the purified RNA was separately subjected to hydrolysis using the Next Magnesium RNA Fragmentation module (-OH) or RNase T1 digestion (T1) (NEB). Reactions were resolved by 8% Urea PAGE, exposed on a phoshorimager screen, and scanned using the Storm 820 imaging system (GE Healthcare). Deduced RNA structures were drawn using the RNA secondary structure visualization tool *forna* (Vienna RNA Web Services).

#### Electrophoretic mobility shift assays (EMSA)

RNA probes used in EMSA experiments were radiolabeled using the protocol described for ribonuclease activity assays. Reactions were incubated at RT for 30 min in buffer containing 20 mM HEPES pH 8.0, 30 mM KCl, 0.01% Tween-20, 0.5 TCEP, 0.2 mg/ml BSA (Sigma-Aldrich), 40 μg/ml of yeast tRNA (Ambion Thermo Fisher), and the indicated amount of purified SOX protein. Reactions volumes were kept at 10 μl and stopped with 3 μl 7x EMSA loading dye (70 mM HEPES pH 8.0, 420 mM KCl, 35% glycerol). Reactions were resolved by 12% native PAGE, and gels were imaged on a Typhoon multivariable imager (GE Healthcare) and quantified using GelQuant software package (Molecular Dynamics).

#### SOX RNA footprinting assay

*LIMD1* 54 RNA was 5’ end labeled with γ-[^32^P]-ATP-6000 Ci/mmol 150mCi/ml (PerkinElmer) using T4 PNK (New England BioLabs). RNA was then gel purified as stated previously. EMSA gel shifts were first used to determine optimal binding conditions (>90% binding, homogeneous complexes of RNA-protein). Binding buffer contained 0.01% Tween 20 (Sigma Aldrich), 5 mM CaCl_2_, 5 mM KCl, 50 mM NaCl, 0.5 mM TCEP, 20 mM HEPES pH 8.0, 0.04 mg/mL yeast tRNA (Ambion), 0.2 mg/mL nuclease free Bovine Serum Albumin (BSA) (Ambion). A dilution series of SOX (8 μM - 0.5 μM) was incubated with 1 μL of radiolabeled *LIMD1* 54 in the presence of 0.1 unit of RNase I (Epicentre Illumina). Reactions were incubated at RT for a total of 10 min before being ethanol precipitated. RNA pellets were then resuspended in 5 μL of 95% formamide solution containing 10 mM EDTA and boiled for 5 minutes. Samples were then loaded onto a 10% analytical grade UREA-PAGE gel and run at 22 watts for 1.5 hours. Gels were imagined and analyzed as stated above. In order to produce an RNase T1 ladder, 1 μL of *LIMD* 54 was incubated with 0.1 units of RNase T1. Reactions were incubated at RT for 5 minutes before being quenched and prepared as stated previously. The *LIMD1* 54 hydrolysis ladder was generated as stated in the in-line probing methods.

#### Bio-layer interferometry (BLI) real-time binding kinetics

RNA probes (3’ end labeled with biotin) were synthesized from Dharmacon (GE healthcare) and HPLC and PAGE purified (See Table S2). The Octet RED96 Bio-Layer Interferometry instrument and Streptavidin (SA) Biosensors were available from ForteBio (Menlo Park, CA). All steps were performed in reaction buffer similar to EMSA binding conditions. Biosensors were incubated with 200 nM of the biotinylated RNA substrate for 180 sec and free RNA was washed away in EMSA buffer containing no RNA. SOX (300 nM) was incubated with the RNA conjugated biosensors for 200 sec, then complexes were dissociated for minimum of 15 min. Response curves for each biosensor were normalized against biosensors conjugated to RNA in the absence of SOX (buffer only control). Normalized response curves were processed using Octet Software version 7 to obtain kinetic values.

**Fig. S1.**
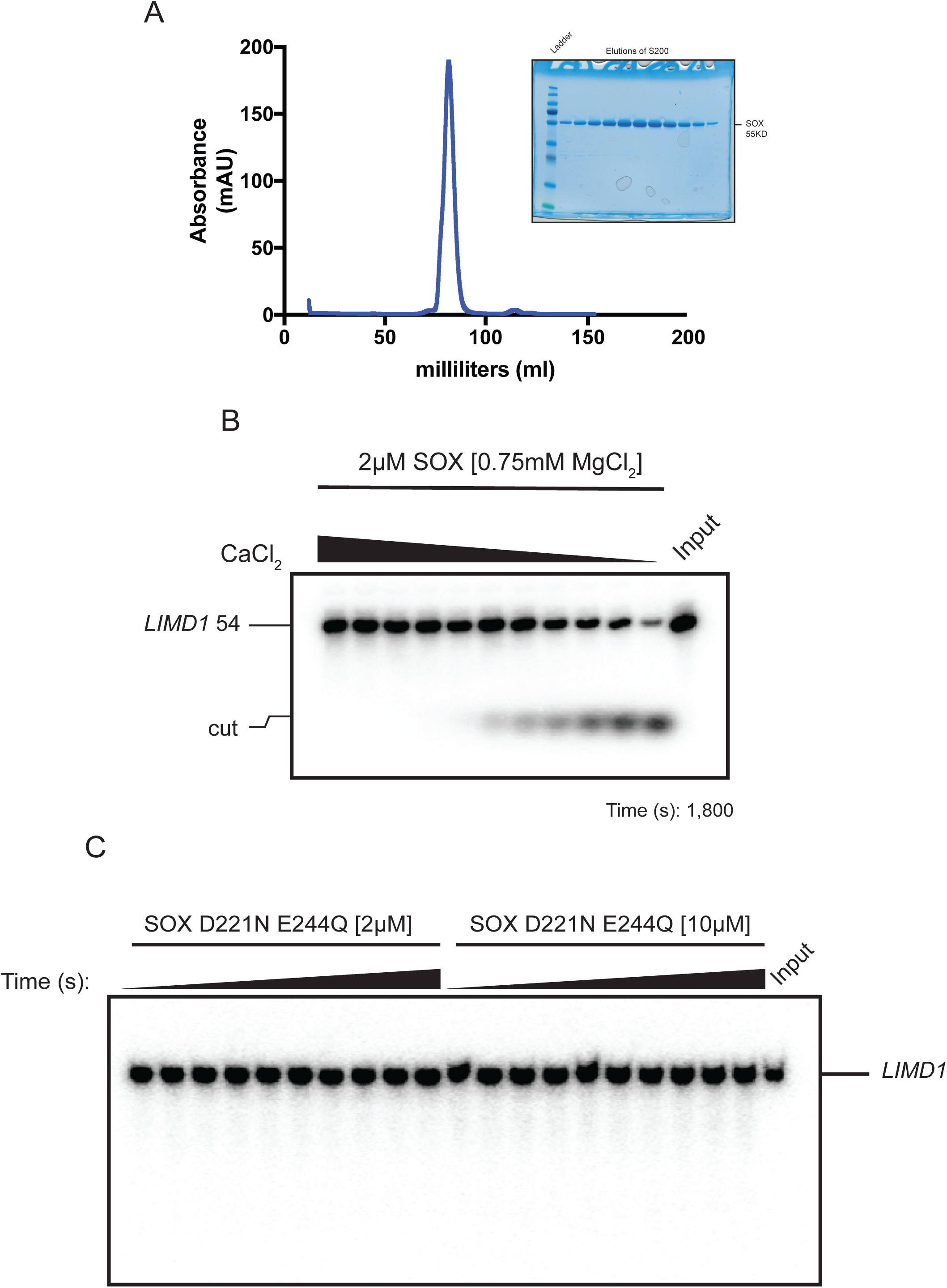
(A) Chromatograph of an S200 size exclusion run of KSHV SOX. SOX eluted predominately as a monomer as determined by molecular weight standards. Inset shows a Coomassie stained SDS-PAGE gel of the >95 pure SOX protein. (B) Raw data used to generate the curve in Fig. 1C. Urea-PAGE gel showing the effect of adding increasing concentrations of CaCl_2_ to reactions containing 2 μM of SOX in reaction buffer containing 0.7 mM MgCl_2_ and 5’-^32^P-labeled *LIMD1 54* RNA. Input lane indicates reaction conditions with 5’-^32^P *LIMD1 54* alone. (C) A time course using an active site mutant of SOX (D221N/E244Q) in the presence of 5’-^32^P *LIMD1 54* over the course of 1 h at low (2 μM) and high (10 μM) concentrations of SOX. Reactions were quenched every 6 minutes.

**Fig. S2.**
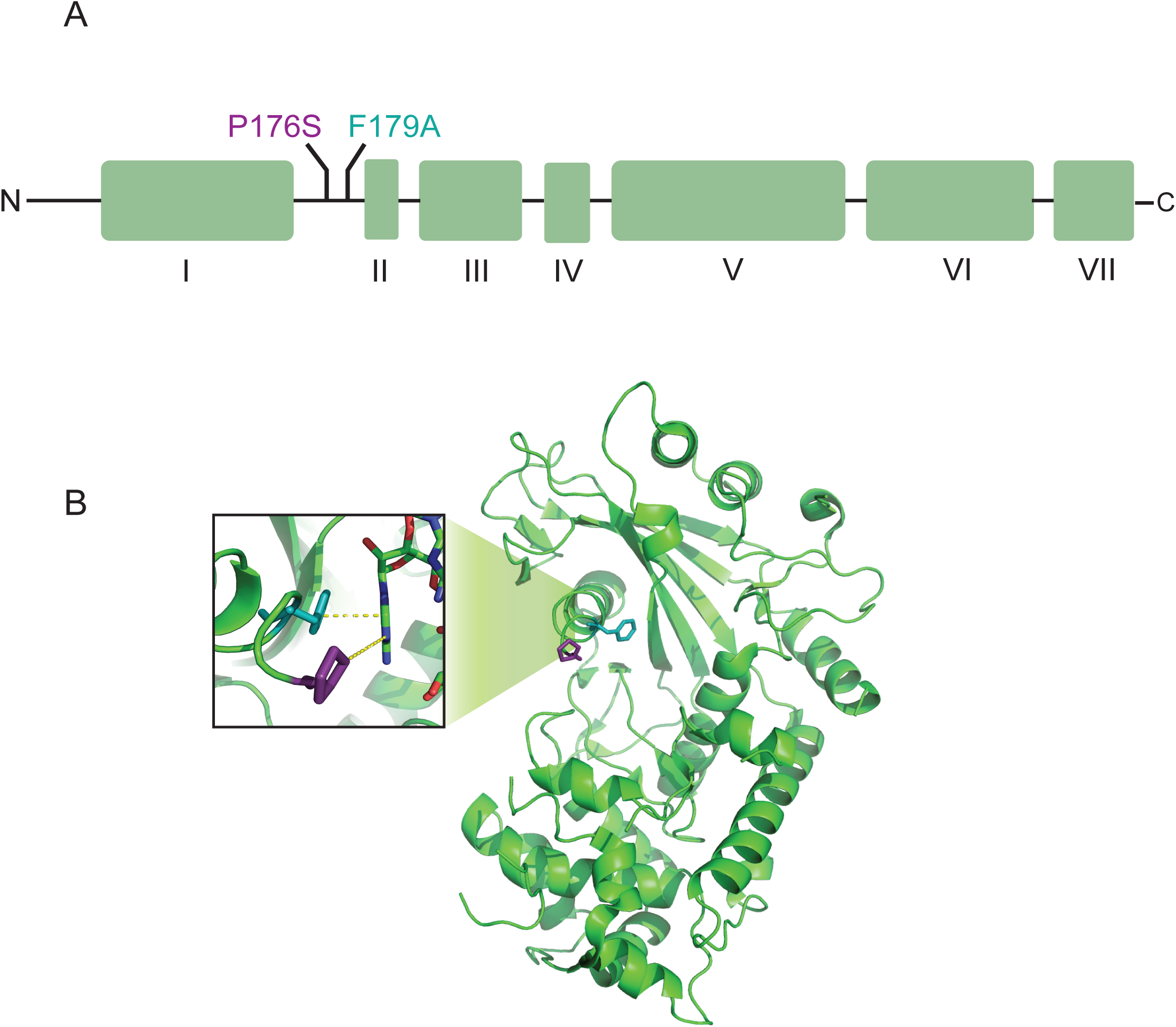
(A) Domain architecture of SOX and locations of the host shutoff mutants. (B) 3D structure and position of SOX host shutoff mutants (PDB ID: 5HSW). Residues P176 and F179 are located within the bridge region of SOX close to the active site. Mutations P179S and F179A are thought to disrupt RNA binding; RNA substrate is hidden for clarity. Inset shows that residues P176 and F179 coordinate an adenine of the RNA substrate.

**Fig. S3.**
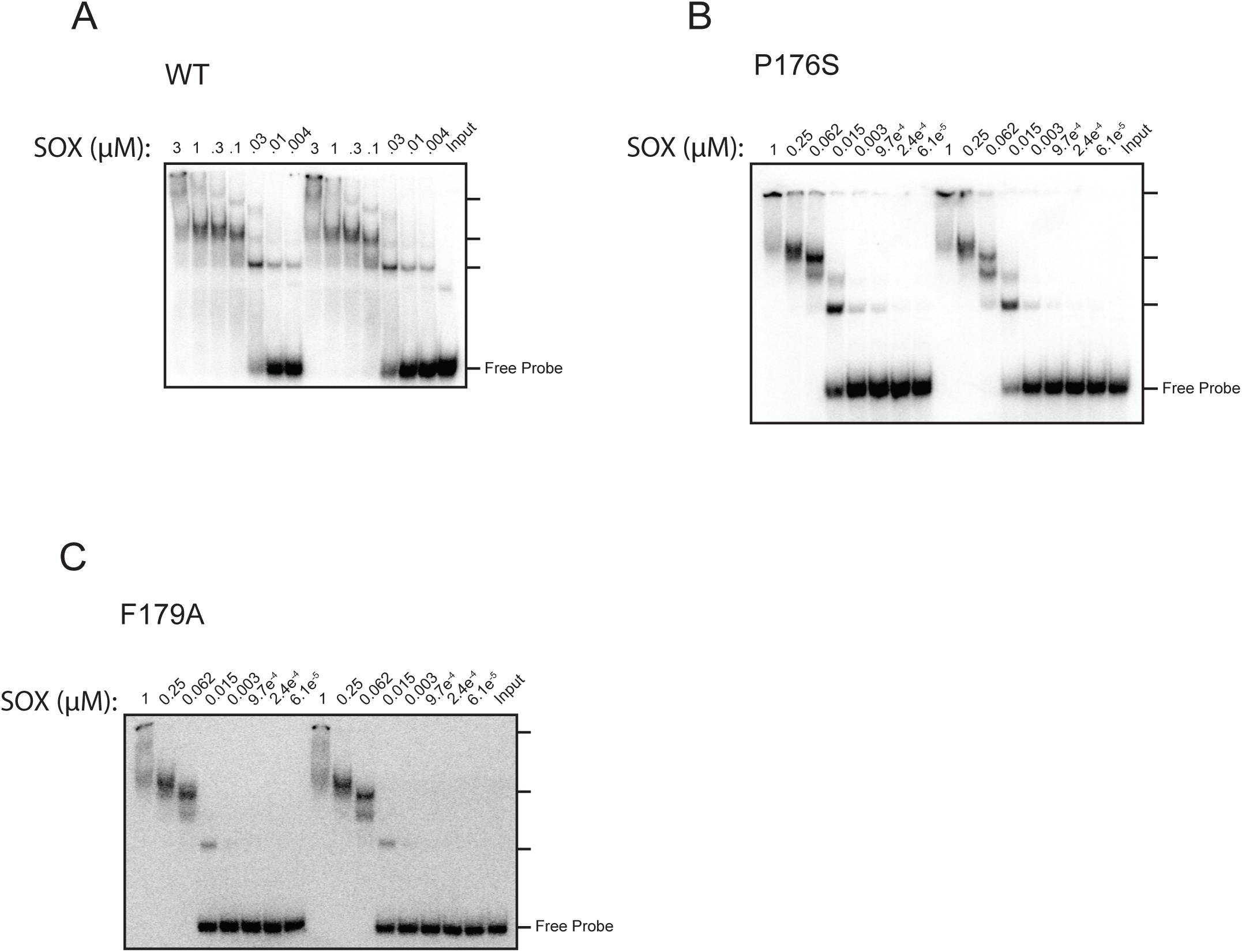
Electrophoretic mobility shift assays (EMSA) of WT (A), P176S (B), or F179A (C) SOX using a 5’ ^32^P-labeled *LIMD1 54* RNA probe. Each reaction was run in triplicate using a 5% native PAGE gel. Gels were dried and quantified by taking the ratio of bound to unbound *LIMD1 54.*

**Fig. S4.**
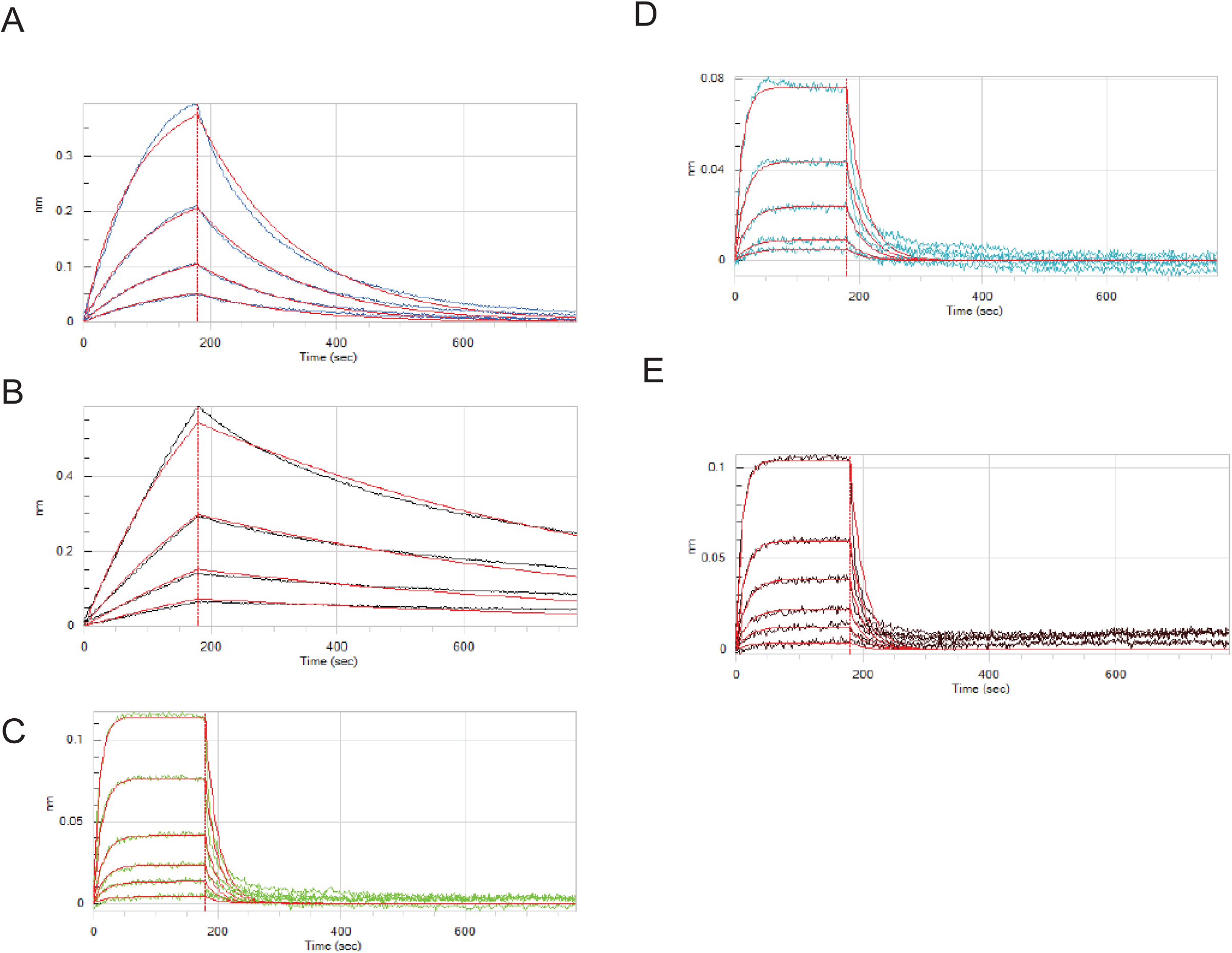
Kinetic characterization of SOX binding to WT *LIMD1* 54 (A), the *LIMD1* 54 mutants *LIMD1 A-G cut site* (B), *LIMD1 3xA-G* (C), and *LIMD1 Zipper* (D), and the miRNA precursor *K2–31* (E). All RNAs were 3’ end labeled with biotin and conjugated to avidin-coated biosensors. Biosensors were then incubated in a well containing SOX to determine on-rates before being transferred to a well containing buffer only to determine off-rates. Red curves represent the statistical fitting of each curve, while the black and colored lines show experimental data.

**Table S1.** Statistical analysis of the Bio-layer interferometry (BLI) data from Figs. S4A-E. All samples were analyzed the same day using BLI-Octet system.

| Load sample | $K_d$ (M) | Full $X^2$ | Full $R^2$ |
| --- | --- | --- | --- |
| *LIMD1* 54 | 1.63E-08 | 0.0572 | 0.9807 |
| *LIMD1A_G* 54 | 2.52E-08 | 0.3251 | 0.9815 |
| *LIMD1* 3xA_G | 71.08E-08 | 0.0292 | 0.9761 |
| *LIMD1* Zipper | 90.40E-08 | 0.1646 | 0.9942 |
| *K2-31* | 10.94E-07 | 0.1508 | 0.9926 |

**Table S2.**
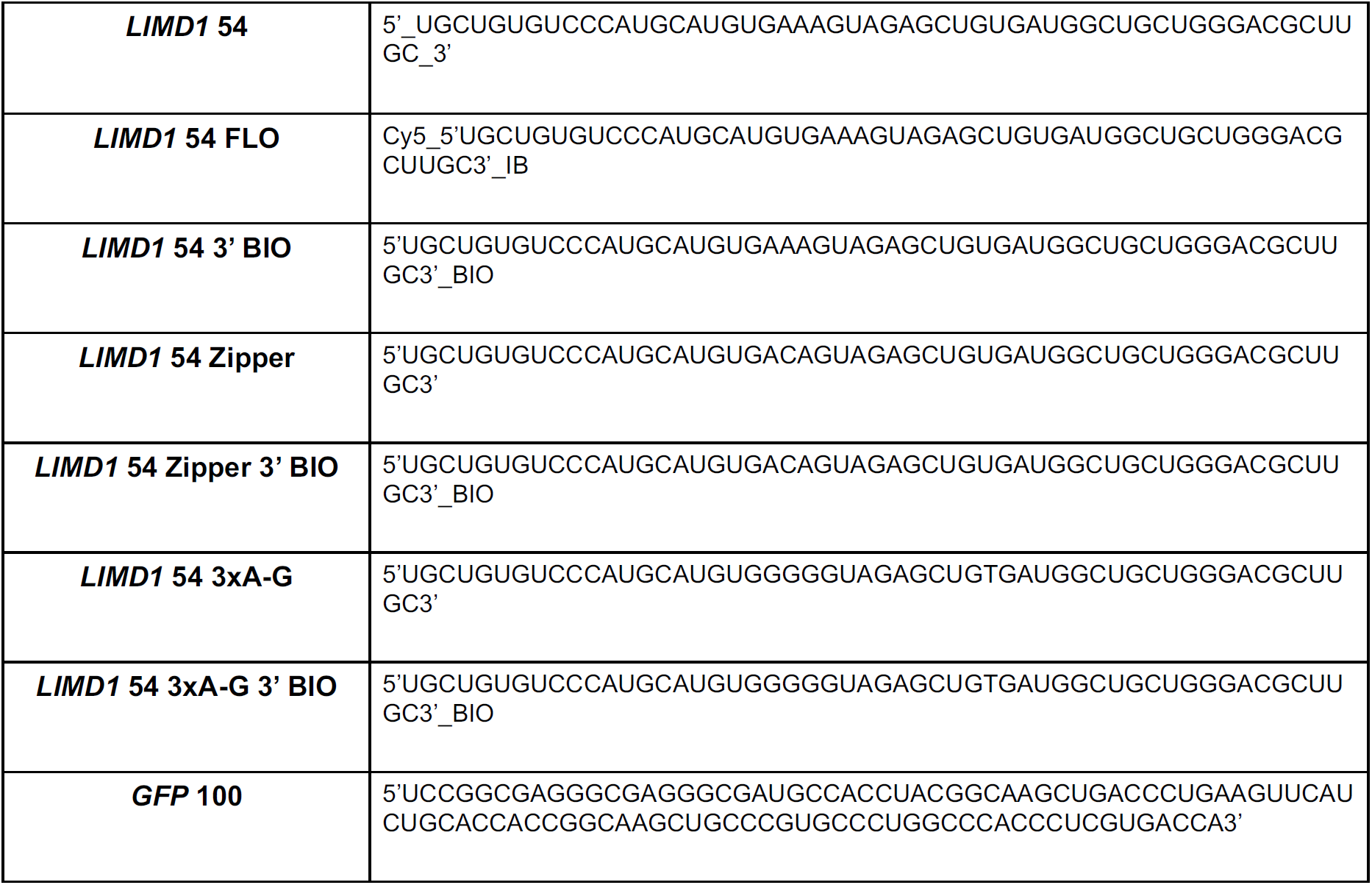
Nucleotide sequences of the RNA substrates used in the binding and turnover assays. Modifications made to either the 3’ or 5’ ends of the RNA are labeled accordingly.

**Fig. S5.**
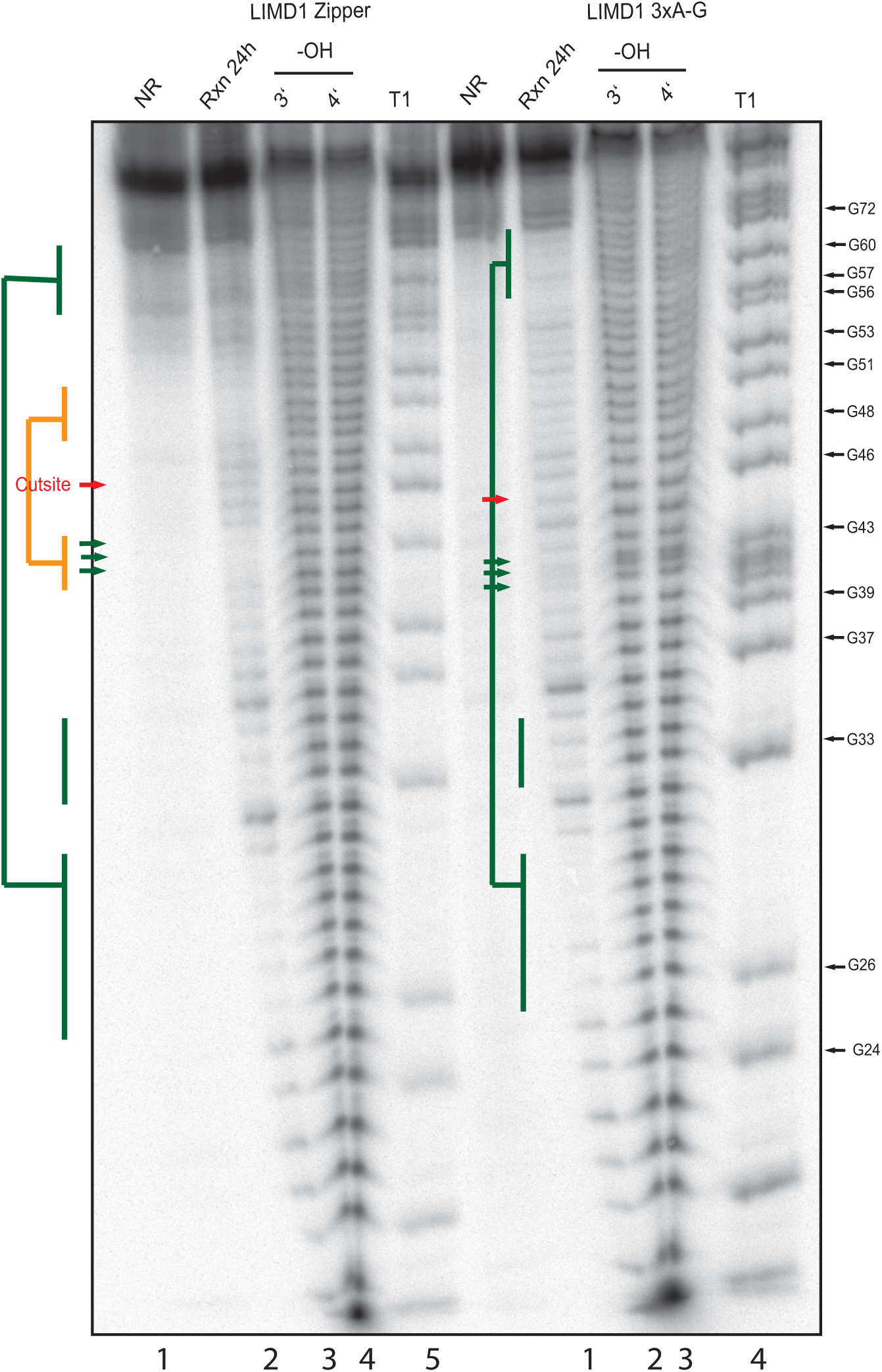
In line probing of the *LIMD1 54* Zipper and 3xA-G mutants to determine the secondary structures shown in Fig. 5A. Lanes 1–5 represent the in-line probing analysis for the LIMD1 Zipper mutant (AAA ➔ ACA) and lanes 6–10 show the analysis for LIMD1 3xA-G. The RNA was loaded directly (NR, no reaction, lanes 1, 6), subjected to cleavage by RNase T1 (lanes 5, 10) or alkaline hydrolysis (-OH, lanes 3, 4, 8, 9), or incubated at room temperature for 24 hours (lanes 2, 7) at pH 8.3 (in-line reaction, Rxn). Samples were separated on an 8% urea PAGE analytical gel. Accessible or unpaired regions generate cleavage bands, whereas structured regions remain blank (orange and green colored). The loop region eliminated in the Zipper mutant is marked by green arrows.

**Fig. S6.**
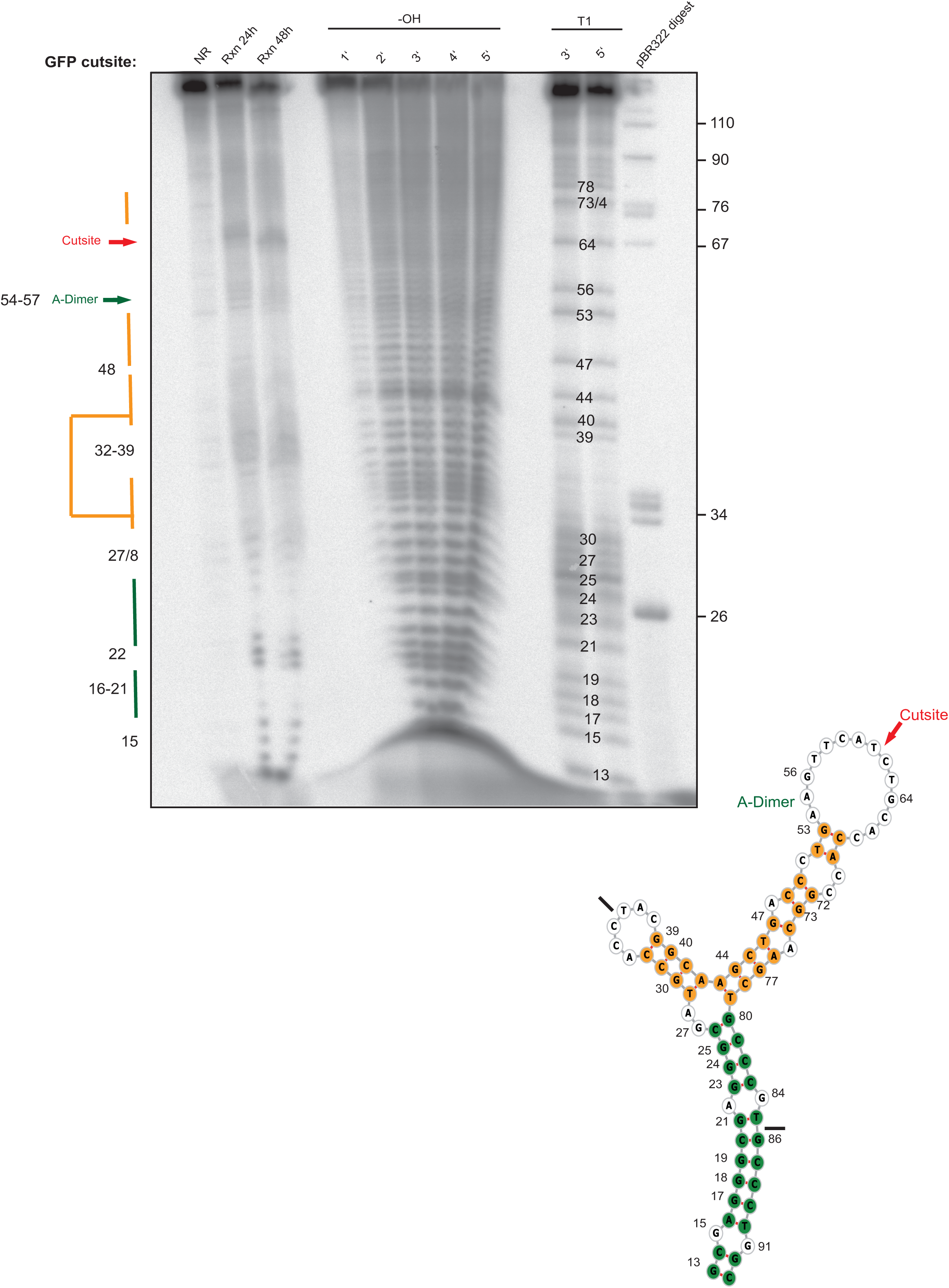
In-line probing analysis of the *GFP* 100 RNA. The RNA was loaded directly (NR, no reaction), subjected to cleavage by RNase T1 or alkaline hydrolysis (-OH), or incubated at room temperature for 24 h or 48 h at pH 8.3 (in-line reaction, Rxn). Samples were separated on an 8 % urea PAGE gel. Accessible or unpaired regions show cleavage bands whereas structured regions stay blank (purple and green colored). The *GFP* 100 RNA structure deduced from the probing gel is shown in the lower right.

